# Genomics of experimental diversification of *Pseudomonas aeruginosa* in cystic fibrosis lung-like conditions

**DOI:** 10.1101/2020.04.14.041954

**Authors:** Alana Schick, Rees Kassen

**Affiliations:** Biology Department and Centre for Advanced Research in Environmental Genomics, University of Ottawa, Ottawa, Ontario, K1N 6N5, Canada

## Abstract

*Pseudomonas aeruginosa* is among the most problematic opportunistic pathogens for adults with cystic fibrosis (CF), causing repeated and resilient infections in the lung and surrounding airways. Evidence suggests that long-term infections are associated with diversification into specialized types but the underlying cause of that diversification and the effect it has on the persistence of infections remains poorly understood. Here, we use evolve and resequence experiments to investigate the genetic changes accompanying rapid, *de novo* phenotypic diversification in lab environments designed to mimic two aspects of human lung ecology: spatial structure and complex nutritional content. After ∼220 generations of evolution, we find extensive genetic variation present in all environments, including those that most closely resemble the CF lung, attributable to a combination of high mutation supply rates resulting from large population sizes and the complex ecological conditions imposed by resource complexity and spatial structure. We use the abundance and frequency of nonsynonymous and synonymous mutations to estimate the ratio of mutations that are selectively neutral (hitchhikers) to those that are under selection (drivers). A significantly lower proportion of driver mutations in spatially structured populations suggests that reduced dispersal generates subpopulations with reduced effective population size, decreasing the supply of beneficial mutations and causing more divergent evolutionary trajectories. The genes most commonly mutated tend to impact regulatory functions linked to a range of CF-associated phenotypes, including one gene that confers antibiotic resistance despite the absence of antibiotic selection in our experiment, but do not appear to be specific to CF-like conditions arising from antimicrobial treatment, immune system suppression, or competition from other microbial species. Our results are consistent with models of adaptation that see the first mutations fixed during adaptation to a stressful environment being those that are broadly beneficial across a range of environments.

## Introduction

Of the many complications associated with the genetic disorder cystic fibrosis (CF), arguably the most difficult to manage and treat is chronic infection of the CF airways by the opportunistic pathogen *Pseudomonas aeruginosa*. Chronic infection occurs in 60-70% of adult CF patients in Canada and is associated with increased morbidity and mortality, irrespective of lung function (Rajan and Saiman 2002; Schaedel et al 2012; Hoiby and Pressler 2006). The majority of infections are thought to result from colonization by environmental strains that adapt to the stressful conditions of the CF lung, leading to characteristic phenotypic and genetic changes including the loss of motility and virulence, a tendency to form mucoid colonies, and the acquisition of high levels of antibiotic resistance (Poole 2005; Smith et al 2006; Mowat et al 2011). The transition to chronic infection is also marked – and, indeed, may be caused – by rapid diversification of *P. aeruginosa* into phenotypically and genetically distinct clones, some of which coexist for years within the same host (Foweraker et al 2005; Ashish et al 2013; Workentine et al 2013; Wright et al 2013; Markussen et al 2014). The causes of diversification, and the contribution of this diversity to the long-term persistence of *P. aeruginosa* infections, remain poorly understood.

Two features of the ecology of microbial life in the CF lung have been shown to promote phenotypic diversification that recapitulates the phenotypic variation seen among clinical isolates from CF patients (Schick and Kassen 2018). The first is the nutritional complexity of the airways and the second is the reduced dispersal among subpopulations caused by both the thick mucous layer covering the lung epithelia and the spatially compartmentalized nature of the lung. Our previous work showed that nutritional complexity was the main driver of within-population diversification, a result consistent with the view that ecological opportunity, the range of underutilized resources, generates strong divergent selection that can drive diversification (Futuyma 1988; Schluter 2000; Kassen 2009). Substantial phenotypic divergence among populations was observed as well, suggesting that reduced dispersal associated with spatial segregation allows distinct populations to explore divergent evolutionary routes to adaptation (Markussen 2014; Flynn et al 2016; Schick and Kassen 2018). The rapid and repeated diversification of *Pseudomonas aeruginosa* demonstrated both empirically and experimentally suggest that this diversification, by generating extensive phenotypic and, presumably, genetic variation on which selection can act, may be a key first step in the development of chronic infections.

Two properties governing the genetics of diversification remain unclear. The first is the extent to which phenotypic variation is matched by comparable levels of genetic variation. Theory suggests that there should be a close correspondence between the two, for three reasons. First, mutation and genetic drift, the fluctuations in allele frequencies associated with finite population size, are stochastic processes that cause replicate populations descended from a common ancestor to diverge through time even in a common environment. Second, strong divergent selection generated by ecological opportunity is expected to lead to the evolution of genetically distinct niche specialists (Schick et al 2015) that can co-exist for prolonged periods of time or even indefinitely (Poltak and Cooper 2011; Traverse et al 2013; Leale and Kassen 2018; Behringer et al 2018). Third, the reduced dispersal associated with spatial segregation allows independently arising beneficial mutations in distinct subpopulations to persist longer than in a well-mixed system because competition among distinct beneficial mutations for fixation arising from clonal interference is weaker. Genetic divergence should thus be closely linked to phenotypic disparity, being most pronounced in ecologically complex environments characterized by abundant ecological opportunity and spatial structure, precisely the conditions thought to characterize the CF airway.

The second is the repeatability of the genetic changes associated with divergence. The probability of parallel evolution, the repeated evolution of the same genetic changes in independently evolved populations under directional selection, is largely a function of population size (Bailey et al 2017): in large populations rare, large-effect beneficial mutations are both more abundant and likely to outcompete independently arising beneficial mutations of smaller effect. For a given population size, however, the probability of parallelism under divergent selection should be lower than under directional selection due to the evolution of genetically distinct niche specialists (Kassen 2014; Turner et al 2018). Spatial structure should also decrease the probability of parallelism relative to a well-mixed system because it reduces effective population size by creating sub-populations that evolve more or less independently of each other. Within patients, then, parallelism is expected to be low.

To test these predictions, we sequenced end-point populations from our previous experiment where *P. aeruginosa* strain Pa14 was allowed to evolve and diversify for ∼220 generations in conditions designed to mimic those encountered during colonization of the CF lung (Schick and Kassen 2018). Briefly, the environments varied in how closely they resemble the resource profile and viscosity of the CF lung, the most CF lung-like environment being composed of a synthetic CF medium (SCFM) based on the chemical contents of samples of CF sputum and mucin, the major protein component of mucus (see Schick and Kassen 2018 for details). The least CF lung-like environment was composed of a minimal medium with glucose as the sole carbon source (MIN) and no mucin. The two intermediate environments were SCFM without the addition of mucin and MIN with mucin, completing the factorial design. We sequenced 81 evolved populations from this experiment to a mean of 120-fold coverage across the genome to uncover all genetic changes that arose and spread to a detectable frequency. These *in vitro* experiments by no means capture all the dimensions of life in the CF lung, as they lack key features such as a diverse microbial community that accompanies *P. aeruginosa* infections and an active immune system. Nevertheless, our results provide insight into the spectrum and quantity of genetic changes accompanying diversification associated with nutrient complexity and spatial structure, two features that are thought to play important roles in governing patho-adaptive dynamics in *P. aeruginosa* during the early stages of colonization.

## Results and Discussion

Comparing whole-genome sequence data from the 81 evolved populations against the ancestral Pa14 genome revealed a total of 656 unique genetic variants, corresponding to an average of 8.10 variants per population. The identity and genomic location of mutations recovered in our analysis are summarized in Figure 1 and 2. The majority of variants were present at low frequencies (mean frequency = 0.207 and median frequency = 0.077), with only 35 variants fixed (i.e., variant frequency = 1). Variants were distributed across the genome (Fig. 1), with peaks corresponding to genomic positions containing genes that are often mutated in *P. aeruginosa* isolates from CF patients (*mexT, lasR*; discussed in more detail below). Nonsynonymous SNPs (single-nucleotide polymorhpisms) were the most common variant class in the experiment, being nearly 3 times more common than the next most abundant class, small indels (Fig. 2A), and there was little variation among environments in the distribution of variant classes (Fig. 2B). This last result suggests that, at this level of categorization at least, there is little genomic signature to distinguish among any of the environmental conditions in our experiment. This interpretation does not hold for the total and average number of variants segregating or fixed among environments, which varied substantially across treatments due to fewer variants being recovered in MIN relative to the other three environments (Fig. 2B; Supplementary Fig. 1; ANOVA, F_(3, 77)_ = 6.24, p < 0.001).

**Figure 1.**
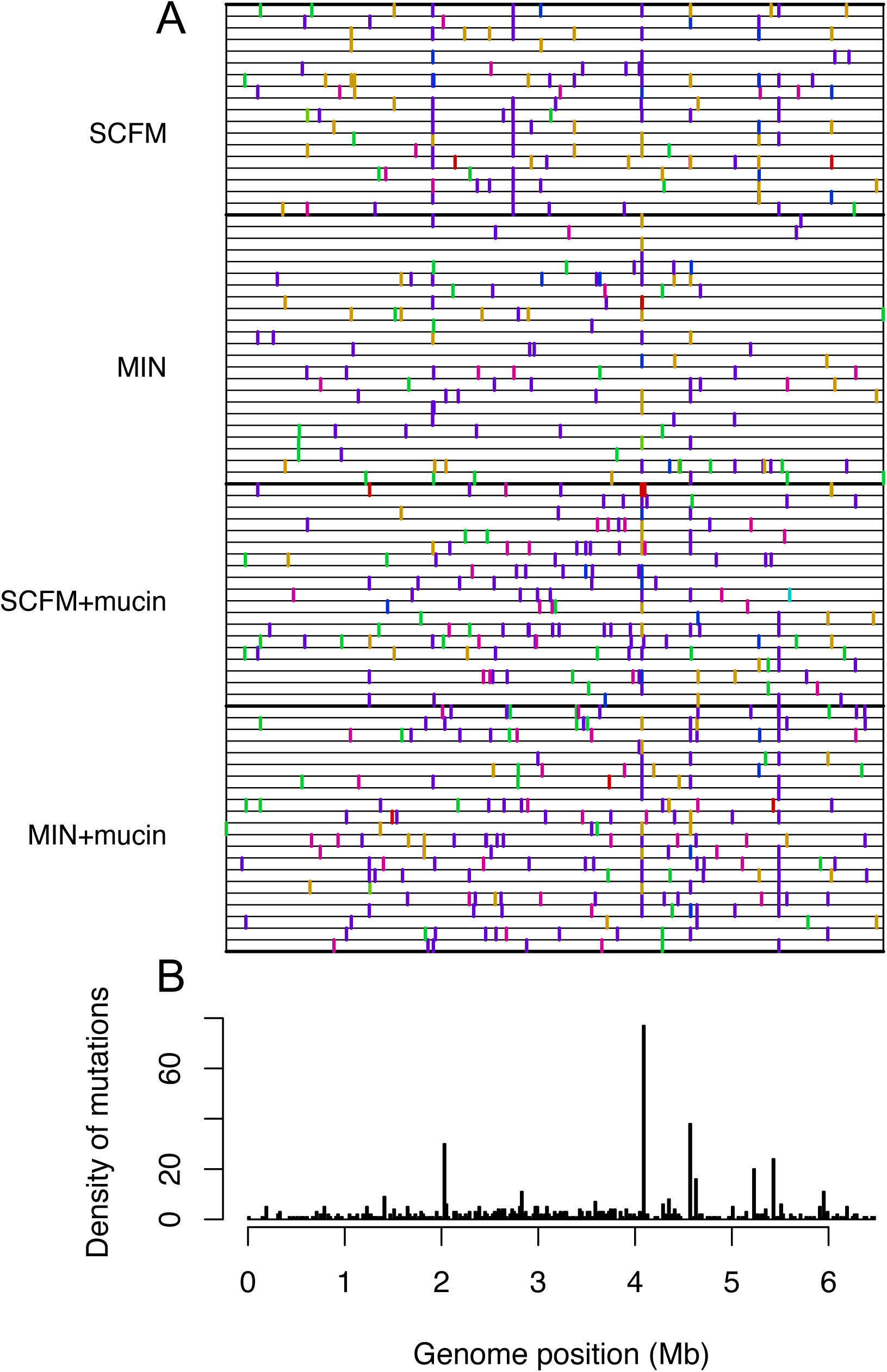
Location of all mutations discovered in evolved populations. **A**. Position in the genome of mutations per population; each row is a population and the genome is represented horizontally. Mutational types are coloured as in Figure 2. **B**. The density of mutations along the genome in 10 kb windows.

**Figure 2.**
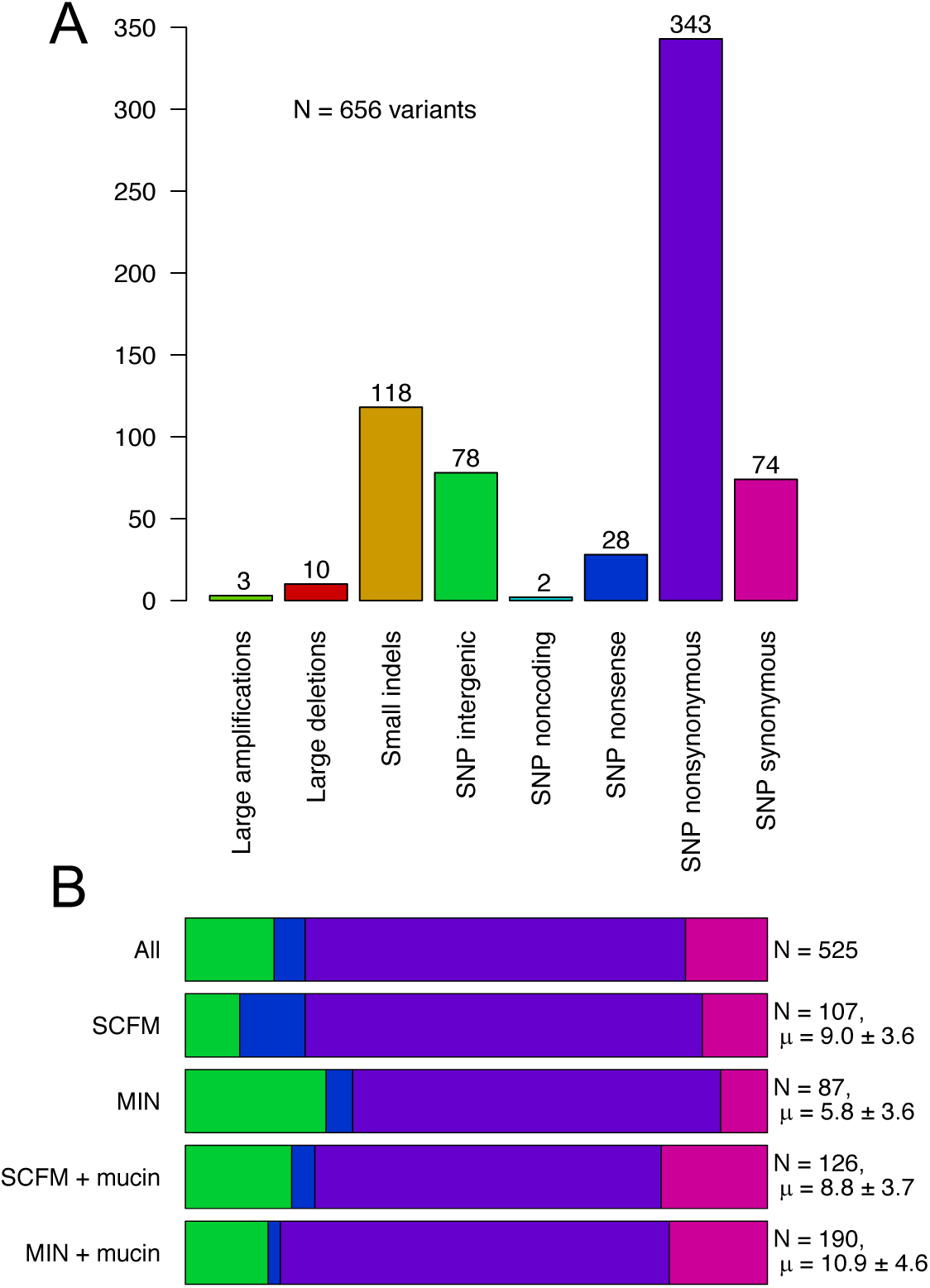
Types of mutations. **A**. The distribution of mutations by mutational type. **B**. The distribution of the different types of only point mutations (SNPs) by selection environment. N is the total number of variants for all populations within each group, with mean and standard deviation of the number of variants per population.

### Positive selection and the prevalence of genomic hitchhiking

To what extent do the patterns of genomic variation we observe in our experiment reflect the action of selection, as opposed to other mechanisms influencing the distribution of genetic variation like the stochastic effects of drift? Selection is likely the dominant driver of genetic variation in our experiment because population sizes were, by design, large (approximately 10^8^ cfu/ml, close to the densities observed in sputum samples of CF patients; Stressman et al 2011) and the experiment relatively short, meaning neutral mutations are highly unlikely to increase in frequency on their own. However, neutral or mildly deleterious variants can reach appreciable frequencies if they occur in the same genetic background as one or more beneficial mutations, a process termed ‘hitchhiking’. The genomic variation we observe in our populations is therefore likely a mixture of beneficial ‘driver’ mutations and hitchhiking mutations; ideally, we would like to know which is which. This task is relatively straightforward when mutations are few in number, as their fitness effects can be evaluated directly through competition experiments (for example Khan et al 2011; Schick et al 2015; Buskirk et al 2017). For studies such as this one, where there are many evolved populations and large amounts of genomic variation within each, direct measurements of fitness on individual variants are impractical and rates of hitchhiking must be inferred using statistical methods.

As a first step, we calculate the ratio of nonsynonymous to synonymous mutations under the assumption that the majority of synonymous mutations are neutral with respect to fitness (although reports to the contrary suggest synonymous mutations can occasionally be adaptive; see Bailey et al 2014; Agashe et al 2016; Kristofich et al 2018; Lebeuf-Taylor et al 2019). Of the 445 SNPs we observed in protein coding regions, 371 and 74 were nonsynonymous and synonymous, respectively, which translates into a frequency of 16.6% synonynmous SNPs. Given that 25.1% of all SNPs in protein coding regions are synonymous in Pa14 (Yang et al 2011), nonsynonymous SNPs are vastly over-represented relative to synonymous SNPs in our experiment, suggesting that positive selection is driving a large number of these polymorphisms to high frequency. Further evidence supporting this inference comes from examining the SNPs present in the five most frequently mutated genes in our experiment (*lasR, morA, mvfR, orfH*, and PA14_32420): all five contain only nonsynonymous changes, suggesting strong positive selection on protein altering mutations in these genes.

Next, we estimate the proportion of driver to hitchhiker mutations by comparing the observed number of nonsynonymous mutations to that expected from the observed number of synonymous mutations, again under the assumption that the latter are neutral. The rationale here is that if nonsynonymous hitchhiker mutations are also neutral, this class of mutation should be at least as frequent as that of synonymous sites in our experiment. The number of putative driver mutations – excluding those beneficial mutations arising late in the experiment that have not had sufficient time to become common – can therefore be calculated by subtracting the number of expected nonsynonymous SNPs under neutrality (hitchhikers) from the number of nonsynonymous SNPs observed. Our results, which are summarized in Table 1, show that 28.4% of all observed SNPs are likely drivers when all populations are considered together, a value that is not too far off the value of 20% estimated for evolving populations of yeast (Buskirk et al 2017). Disaggregating this result by environment reveals a striking result: environments lacking mucin harbour a larger proportion of putative drivers than those with mucin (Table 1; χ^2^ = 7.638, *P* = 0.008). As population sizes and mutation rates are similar across environments, these results are unlikely to reflect lower mutation supply rates in the presence of mucin. If anything, the opposite is the case, as the total number of nonsynonymous and synonymous mutations tends to be higher in the presence of mucin than in its absence. The simplest explanation for this result is that the presence of mucin, by increasing viscosity and reducing dispersal, creates subpopulations with lower effective population sizes (*N*_*e*_) relative to a well-mixed system, decreasing the supply of beneficial mutations to those with selection coefficients greater than ∼ 1/*N*_*e*_.

**Table 1.** Estimated proportion of SNPs that are driver mutations. The estimated number of hitchhiker SNPs is calculated as the observed number of synonymous SNPs times three (following the expected ratio under neutral evolution of 25.1:74.9 of synonymous to nonsynonymous sites). The estimated number of driver mutations is calculated by subtracting the expected number of nonsynonymous hitchhiker SNPs from the observed number of nonsynonymous SNPs. Proportion is determined by dividing number of nonsynonymous driver mutations by the total number of observed SNPs (including SNPs in noncoding regions).

| Treatment | Observed nonsynonymous SNPs | Observed synonymous SNPs | Estimated number of hitchhiker SNPs | Estimated number of driver SNPs | Estimated proportion of driver SNPs (%) |
| --- | --- | --- | --- | --- | --- |
| All | 371 | 74 | 222 | 149 | 28.4 |
| SCFM | 85 | 12 | 36 | 49 | 45.8 |
| MIN | 59 | 7 | 21 | 38 | 43.7 |
| SCFM + mucin | 82 | 22 | 66 | 16 | 12.7 |
| MIN + mucin | 131 | 32 | 96 | 35 | 18.4 |

### Frequency spectra of nonsynonymous and synonymous SNPs

Examining the frequency spectra of nonsynonymous and synonymous SNPs from all populations reveals the distributions to be very different. Most synonymous SNPs were rare, 97% having a frequency of 0.16 or less, whereas nonsynonymous SNPs were present at a range of frequencies from rare to fixed (Fig. 3). The paucity of high frequency synonymous SNPs suggests that the majority of this class of mutation do not contribute to adaptation, a result that is not surprising in light of the widely accepted view that the majority of synonymous mutations are neutral with respect to fitness. Two notable exceptions were synonymous SNPs in PA14_49020 and gdhB, an NAD-dependent glutamate dehydrogenase, that achieved frequencies of 94% and 36%, respectively, suggesting these mutations could be adaptive. These results notwithstanding, the majority of synonymous SNPs are likely hitchhikers whose frequencies, by definition, must be equal to or less than those of their respective driver mutations. If the same is true for rare nonsynonymous SNPs, then a conservative estimate (because it ignores potentially rare beneficial nonsynonymous mutations) of the frequency of driver SNPs is 31.2 %. Reassuringly, this estimate is not far off that derived above based on inferences based on the ratio of nonsynonymous to synonymous mutations, independent of mutant frequency.

**Figure 3.**
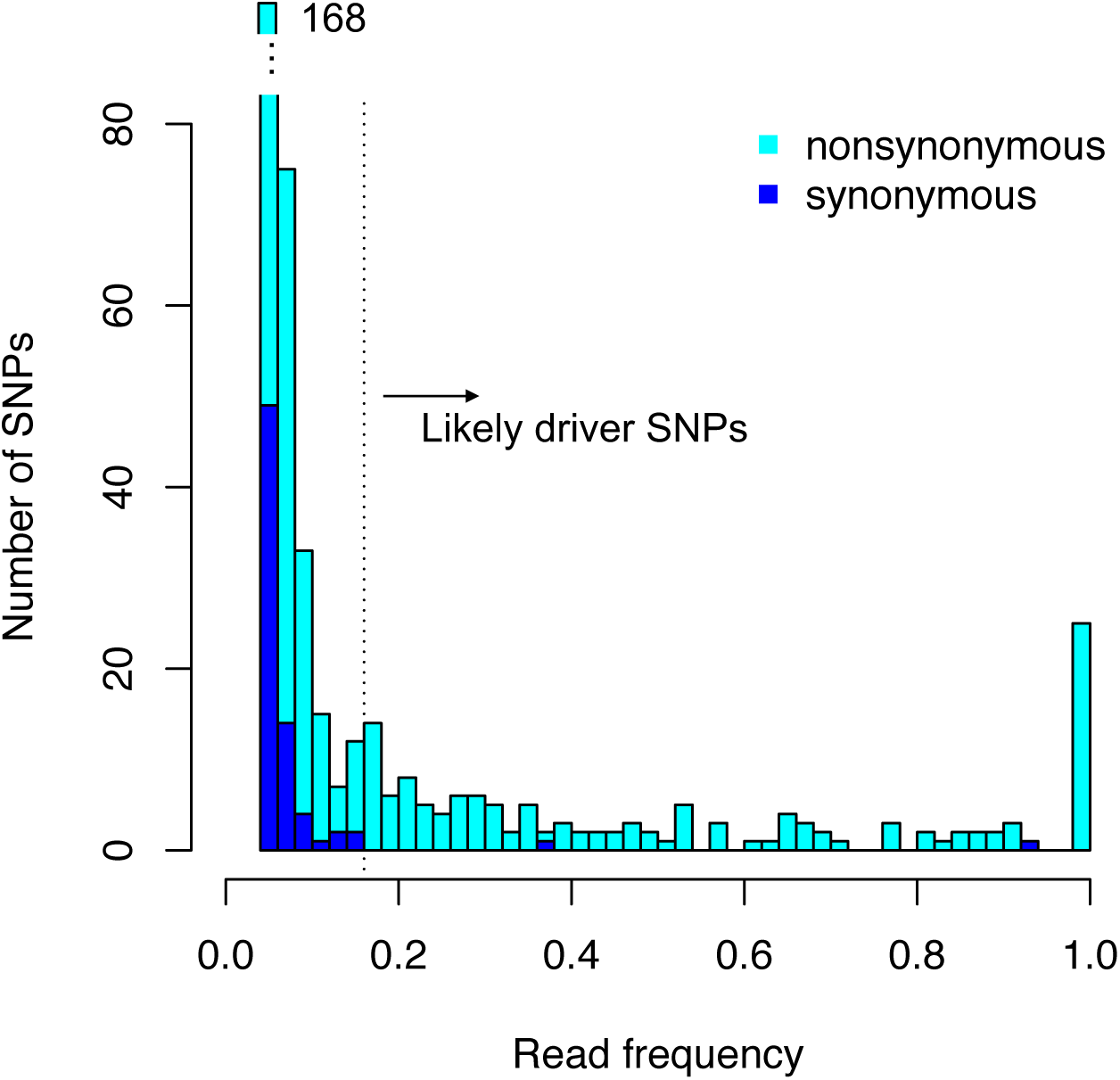
Frequency spectra for nonsynonymous and synonymous SNPs. Distribution of read frequencies for all SNPs found across all evolved populations, nonsynonymous (light blue) and synonymous (dark blue). The majority of nonsynonymous SNPs present at frequencies greater than 0.16 (the range of frequencies of most synonymous SNPs) are likely to be driver mutations.

### The correlation between genetic diversity and phenotypic disparity

To what extent is the phenotypic disparity we observed in our previous work (Schick and Kassen 2018) underlain by comparable levels of genetic diversity? To answer this question we first calculated mean heterozygosity within each population after excluding synonymous and low-frequency (<0.16) nonsynonymous SNPs (as defined above) and then regressed these values against the extent of phenotypic disparity, calculated as the multivariate Euclidean distance among pairs of isolates from the same population (details provided in Schick and Kassen 2018). Average heterozygosity was significantly greater than zero for all treatments (*t*-test, *P* < 0.0001 for all treatments; Fig. 4A), a result that is likely a consequence of high mutation supply rates that generates clonal interference and introduces hitchhiking mutations alongside genetic variants associated with adaptive divergence. As expected from theory, we see a statistically significant positive correlation between heterozygosity and phenotypic disparity among populations within environments (Pearson coefficient = 0.393, *P* = 0.006; Fig. 4B). While SCFM, the condition that resembles the nutritionally complex conditions of the CF lung, had the highest average heterozygosity, as expected, we could not detect an effect of environment on the slope nor the y-intercept of the regression of heterozygosity on phenotypic diversity (ANCOVA, *P* = 0.131 and *P* = 0.630 for treatment and interaction term for treatment and heterozygosity). Together, these results suggest that rapid and extensive phenotypic diversification associated with nutritional complexity is driven by a combination of divergent selection on a few mutations of large effect in the context of a high mutation supply rate. It will thus be difficult to identify the mutations contributing most to divergence from the rich suite of hitchhiking mutations, some of which could modulate the effects of strongly selected mutations, using purely genomic approaches.

**Figure 4.**
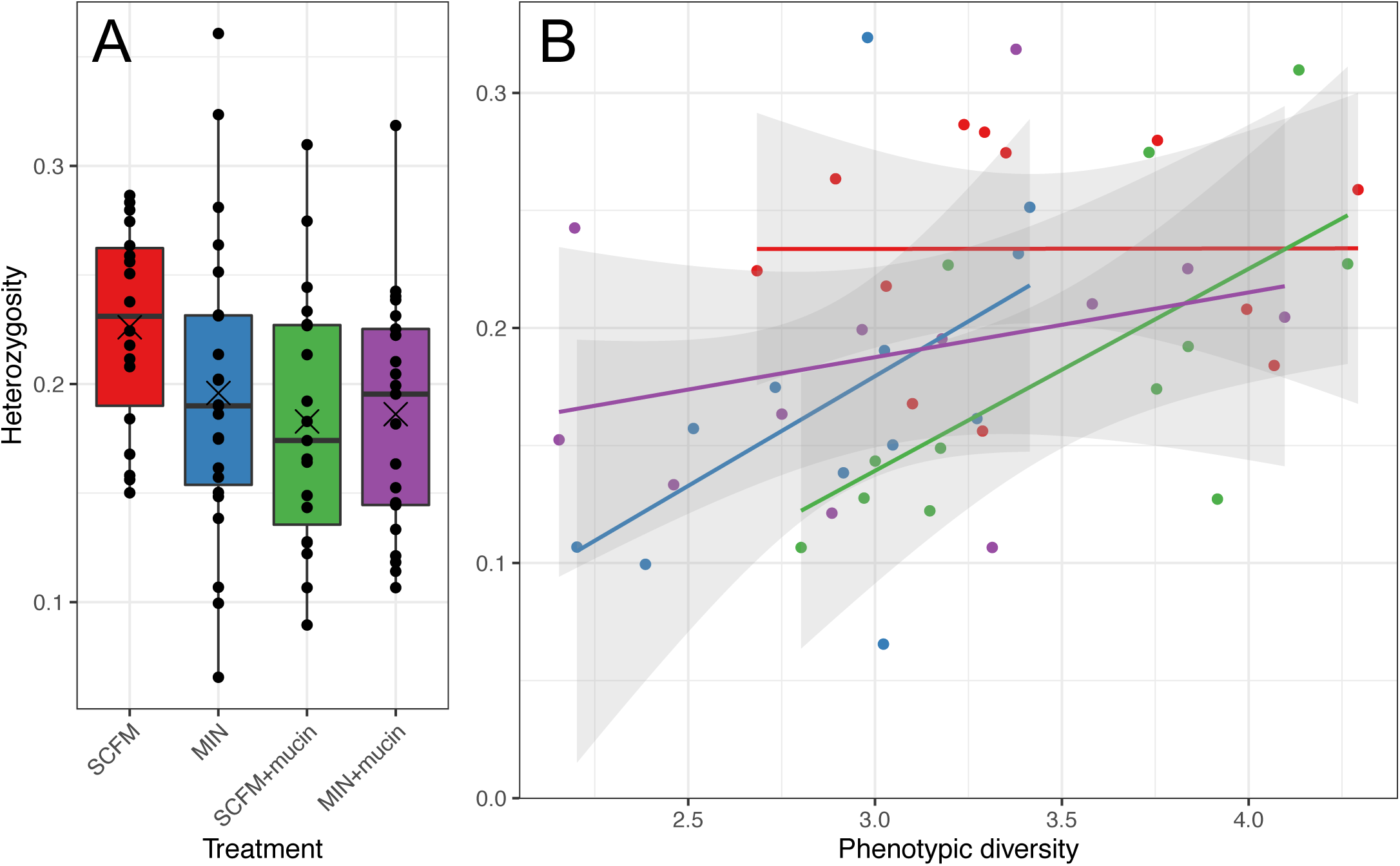
Genetic diversity within populations. **A**. Population mean heterozygosity, calculated using frequencies of variants at all loci with a variant present in a given population, excluding synonymous and low-frequency variants. Horizontal bars represent treatment median and crosses represent treatment mean. **B**. Correlation between heterozygosity and phenotypic diversity (measured by mean euclidean distance between pairs of populations for a suite of phenotypic characteristics, see Schick and Kassen 2018 for details).

### Genome scale patterns of parallelism and constraint

Adaptation to the CF lung typically involves a handful of phenotypic changes involving loss of motility, reduced virulence, increased antibiotic resistance, and mucoidy (Smith et al 2006; Mowat et al 2011). Such repeated, or parallel, evolution of the same phenotypes in independent lineages is often taken to be a marker of strong selection. Whether these parallel phenotypic changes are underlain by parallel genetic changes remains unclear. The consensus from longitudinal (Smith et al 2006, Marvig et al 2015) and cross-sectional (Dettman et al 2013) studies is that, while select genes show high levels of parallelism, genetic parallelism is usually low, there being many genetic routes to putatively patho-adaptive phenotypes. Whether this variation in parallelism is a product solely of the stochastic nature of mutation-driven adaptation or a function of varying degrees of divergent selection favouring different genes in distinct niches of the CF lung is unknown (but see Wong et al 2012, Bailey et al 2015, Turner et al 2018 for more general treatments of this problem). An answer is important because high levels of parallelism that are unique to the CF lung could be used as a genetic marker of the onset of chronic infection. Our experiment allows us to answer this question and test hypotheses about how parallelism is impacted by the environmental complexity associated with the CF lung.

To quantify gene-level parallelism among mutations likely to be under selection, we first restrict our analysis to putative driver mutations by excluding synonymous and low-frequency SNPs that we expect to be hitchhiking, as discussed above. While this approach likely inflates our estimates of parallelism, it should do so in a way that is common across treatments and so should not introduce any systematic bias. Our results are summarized visually in Figure 5 and presented more quantitatively in Table 2. As there is no broadly accepted metric for quantifying gene-level parallelism, we used three distinct measures: variance in dispersion of Euclidean distances between populations, Jaccard index, and observed repeatability relative to expectation under randomness using the hypergeometric distribution (C-score; Yeaman et al 2018). Details on how each metric is calculated are provided in the Methods. For all metrics reported in Table 2, divergence was lowest (and thus parallelism was highest) in the MIN environment and next lowest in SCFM, consistent with the hypothesis that divergent selection reduces the probability of parallelism relative to directional selection. Divergence was highest (parallelism lowest) in environments containing mucin, although the rank-order depends on which metric is used, consistent with the idea that spatial structure and reduced dispersal lead to more divergence among subpopulations within a lineage.

**Table 2.** Parallelism and constraint. Population level patterns of parallelism within each treatment. Dispersion is the mean distance from the centroid of a PCoA (principal coordinates analysis) based on euclidean distance; larger values signify more divergent populations and therefore less parallelism. Jaccard distance is dissimilarity between sets; a Jaccard distance of 1 for any two populations signifies they are completely dissimilar and have no gene variants in common, therefore larger values signify more divergent populations and less parallelism. C-scores measure observed repeatability relative to expectations under randomness; smaller values signify less constraint and therefore less parallelism. For all metrics, SE is standard error and treatments are ranked from most divergent (1) to least divergent (4), equivalent to least parallel (1) to most parallel (4). All three metrics were calculated excluding synonymous and low frequency SNPs. Significance is determined by an ANOVA for Dispersion and by an exact test (permutation test with 10,000 permutations) for Jaccard Index and C-score.

| Measure | Dispersion |  |  | Jaccard Index |  |  | C-score |  |  |
| --- | --- | --- | --- | --- | --- | --- | --- | --- | --- |
|  | Mean | +/- SE | Rank | Mean | +/-SE | Rank | Value | +/-SE | Rank |
| SCFM | 0.710 | 0.054 | 3 | 0.833 | 0.013 | 3 | 4.70 | 0.351 | 3 |
| MIN | 0.457 | 0.055 | 4 | 0.810 | 0.022 | 4 | 4.98 | 0.464 | 4 |
| SCFM + mucin | 0.760 | 0.058 | 2 | 0.856 | 0.020 | 1 | 4.04 | 0.482 | 1 |
| MIN + mucin | 0.855 | 0.063 | 1 | 0.851 | 0.012 | 2 | 4.17 | 0.285 | 2 |
| Significance | $F_{(3,77)} = 4.78; P = 0.004$ | | | Exact test; $P = 0.254$ | | | Exact test; $P = 0.261$ | | |

**Figure 5.**
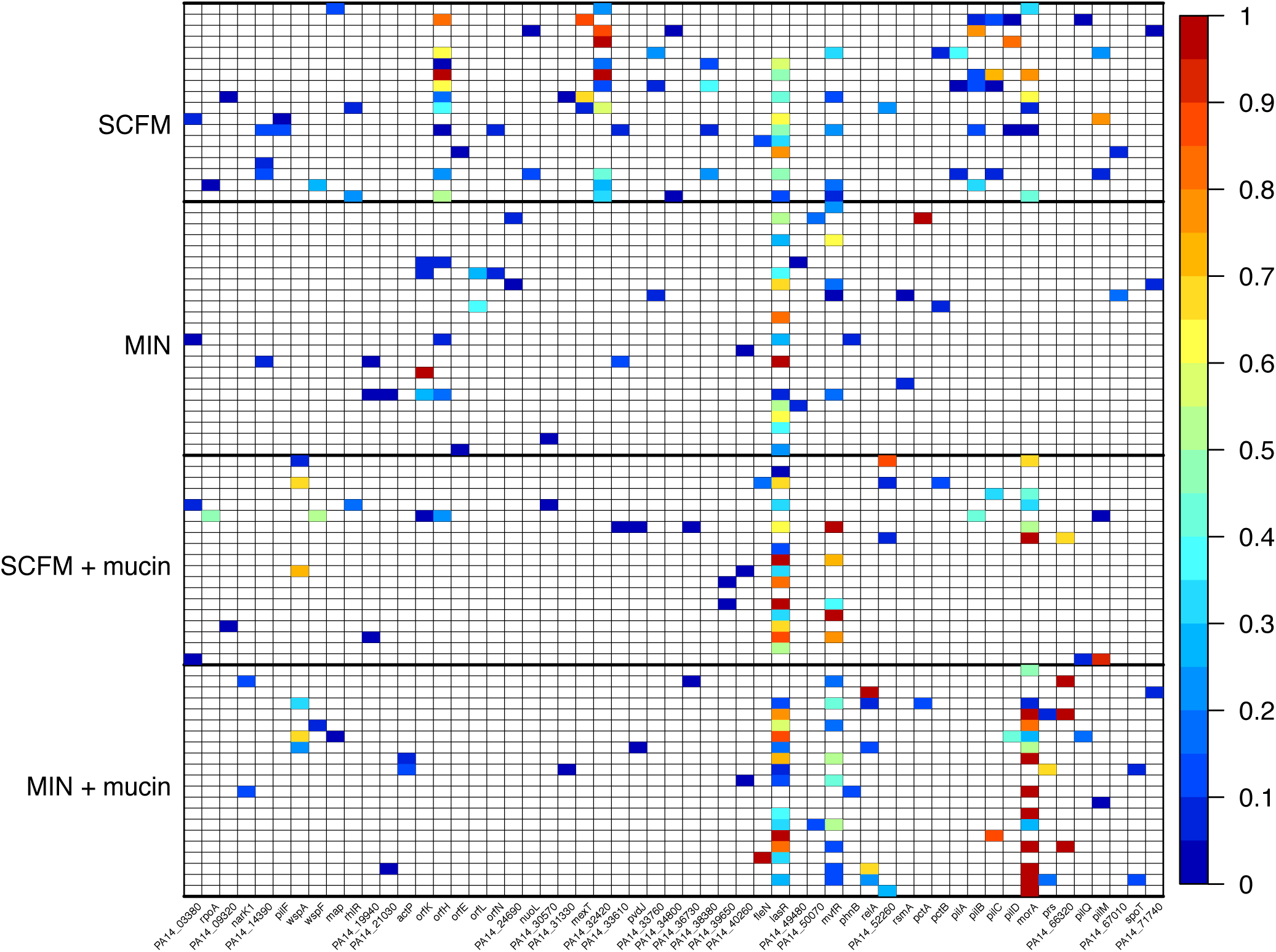
Variant frequencies for multiple use genes. Frequency of variants for genes that have variants in more than one population, after excluding synonymous SNPs (N = 55 genes). Each row represents a population, with the presence of a mutation in each gene indicated by a coloured box (high frequency mutations in red and low frequency mutations in blue). Genes are arranged by genomic position.

### Gene level parallelism and environmental specificity

The spectrum of parallel genetic changes evolved in each environment, shown in Figure 6, reveals two striking features. The first is that, in line with the results from comparative genomic studies (Smith et al 2006, Marvig et al 2015, Dettman et al 2013), most genes show modest levels of parallelism, with repeated changes occurring in just a handful of lines. Notable exceptions are mutations in *lasR* and *mvfR*, both transcriptional regulators involved in quorum sensing, which were shared by the majority of lines across all treatments. The second is that, with a handful of exceptions (*morA* in MIN+mucin or *pilA* and PA14_32420 in SCFM), there is no strong signal of environment-specific parallelism. To quantify this effect and to identify genes with high levels of environment-specific parallelism, we ask whether there is a significantly higher probability of gene-level parallelism, measured as the proportion of populations with non-synonymous mutations in a given gene, within than across treatments. We included only genes that were mutated in more than one population (N = 55 genes) and defined significance using a binomial distribution to calculate the probability of seeing the observed proportion of populations if gene use was random across treatments. A probability less than 0.05 suggests parallelism in that gene is treatment specific. We found 13 genes to be significantly treatment specific in at least one selection environment. The SCFM environment showed the largest number of environmentally specific genes (n= 8) though mutations in three genes (*pilA, pilB*, and *pilC*) likely result in a similar phenotypic effect, namely, a reduction in twitch motility (Burrows 2012). Both the SCFM+mucin and MIN environments were found to have only a single gene mutated in more populations than expected (Pa14_39650 and *orfK*, respectively). Grouping genes by functional classification revealed no patterns, unsurprising given the limited number of treatment specific genes (n =13) and the large number of functional classes. Together, these results suggest that, with the exception of a handful of genes, parallel genetic evolution may be an unreliable marker of CF-specific adaptation.

**Figure 6.**
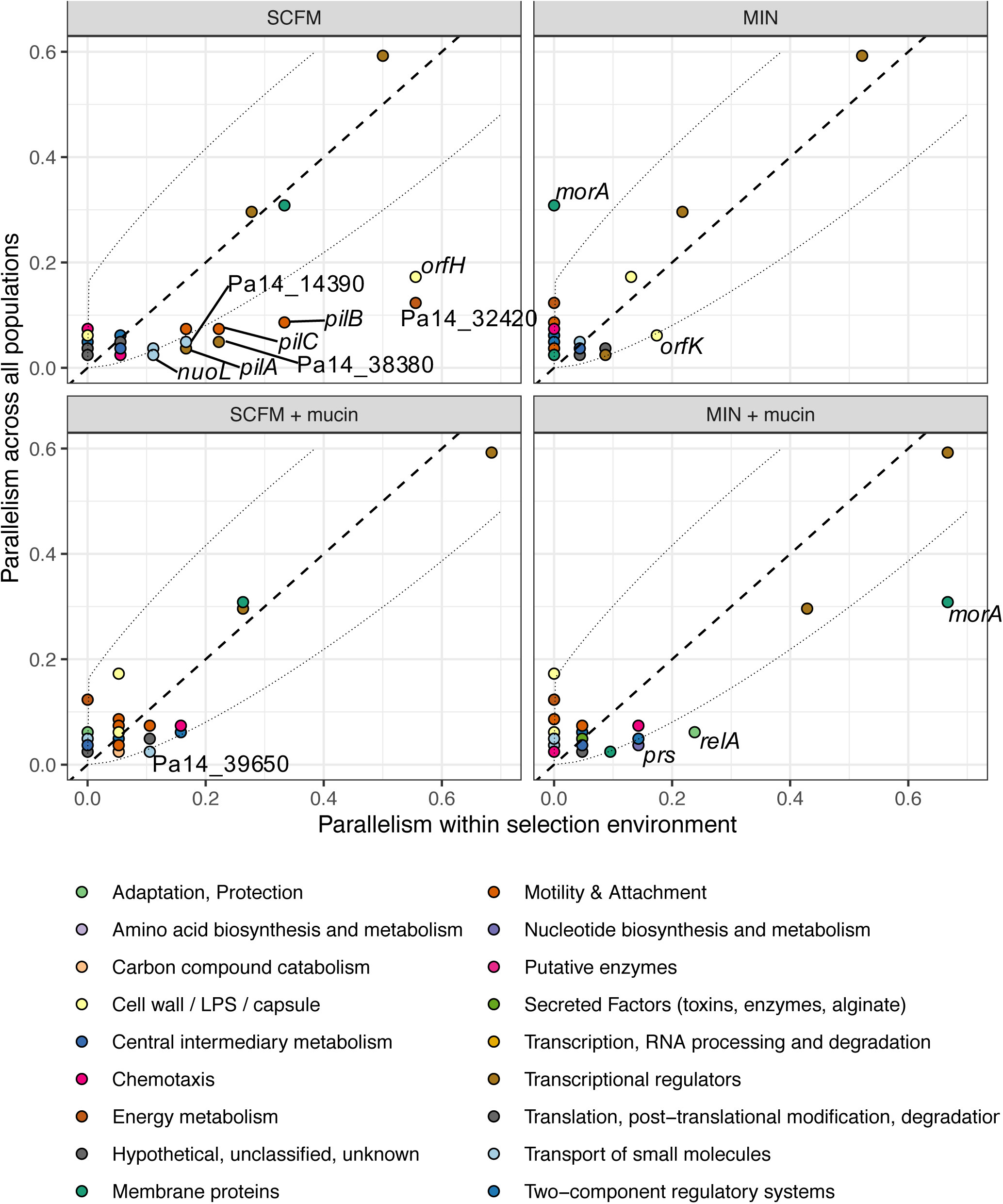
Gene-level parallelism. Parallelism within selection environment versus parallelism across all populations, for all genes with variants in more than one population, after excluding synonymous SNPs. Parallelism for a gene is defined as the proportion of populations with mutations in that gene. Dashed lines represent the line where parallelism within selection environment is equal to parallelism across all populations; points below this line are genes that are more commonly mutated in that environment and points above this line are genes less commonly mutated. Dotted lines represent statistical significance, based on the binomial distribution and an alpha = 0.05. Genes that fall outside the dotted lines are labelled.

### Genomic targets of selection

Our previous work showed that many of the characteristic phenotypic changes that are the hallmark of the onset of chronic infection also evolve under conditions that mimic the nutritional complexity and spatial structure of the CF lung *in vitro*, in the absence of an active immune system and diverse microflora (Schick and Kassen 2018). Is the same true at the genetic level? To some extent, yes. Many of the genes mutated in our experiment are also observed in genomic analyses of isolates from the CF lung, including *lasR, mexT, morA, wspA/F*, and the *pilA-D/Q* genes (Caballero et al 2015, Marvig et al 2015, Freschi et al 2018) and a number serve central roles as regulators of various functions such as quorum sensing (*lasR*; Lafayette et al 2015) small molecule efflux (*mexT*; Kohler et al 1999), and cyclic-di-GMP signaling associated with biofilm formation (*morA, wspF*; Wong et al 2012, Hickman et al 2005; Flynn et al 2016). Few, however, show strong associations with specific components of the CF lung, with the possible exception of *mexT*, which arose multiple times in SCFM, and *morA*, which evolved repeatedly in all environments except MIN. We also failed to observe some characteristic mutations altogether, especially those linked to alginate production such as *mucA, gacA/S*, or *algG/U*, however this is not surprising since we did not observe mucoid colonies in our evolved populations. Our results suggest that many of the genetic changes thought to be characteristic of chronic infections are not specific to the nutritional complexity and viscosity of the CF lung. Rather, many of the genes thought to be hallmarks of adaptation to the CF lung harbor mutations that are beneficial across a range of environmental conditions and serve regulatory roles. Regulatory mutations are often among the first mutations selected in laboratory evolution experiments (Kassen 2014, Dettman et al 2012) and are commonly observed in comparative studies of adaptation to the CF lung (Marvig et al 2015), likely because they help serve to restore the ability to grow under a broad range of stressful conditions. This result is consistent with simple models of adaptation that predict the first mutations fixed during an adaptive walk should not have strongly antagonistic effects across different environments (Martin & Lenormand 2006, Schick et al 2015).

Two additional features of our results deserve mention. The first is the high repeatability of *lasR* mutations, with 74 unique variants being identified in 48 populations in our experiment. Putatively loss-of-function mutations in *lasR* are commonly observed in CF isolates, their selective advantage thought to be the result of reduced expression of acute virulence factors or growth advantages linked to amino acid metabolism (D’Argenio et al 2007, Hoffman et al 2009, LaFayette et al 2015). While we do not know for certain the effect our mutations have on the gene product of *lasR* itself, given that all but one (a large amplification that occurred in the MIN environment) were either nonsynonymous (n = 43), nonsense (n = 5), or small indels (n = 25), it seems likely that our collection includes many loss-of-function mutants. That *lasR* mutations evolved repeatedly in both SCFM and minimal media suggests that growth advantages associated with amino acid metabolism are not responsible for their prevalence *in vivo*. The complexity of quorum-sensing regulation in *P. aeruginosa* can allow mutations in *lasR* to have little or no effect on other quorum-sensing gene products (Feltner et al 2016), suggesting that the fitness advantage of a *lasR* mutants may derive simply from reducing the metabolic cost of expression under high cell densities. This is a hypothesis that awaits further testing.

The second is the prevalence of putatively antibiotic-resistant mutations in our experiment despite the fact that antibiotics were not present at any time in our experiment. Specifically, we observed the evolution of resistance to the fluoroquinolone ciprofloxacin, commonly used to manage *P. aeruginosa* infections in CF patients, in three evolved populations from SCFM (Schick and Kassen 2018). Our genomic analysis points to mutations in *mexT*, a transcriptional regulator of the MexEF-OprN efflux pump, as the likely cause (Breidenstein 2011). Two of the three distinct nonsynonymous SNPs were present at high frequencies of 0.89 and 0.71, suggesting strong selection at this locus, while the third was present at a lower frequency of 0.09. The mechanism by which mutations in this gene confer an antibiotic-independent increase in fitness is unknown, but it could be a response to disulfide stress (Fargier et al 2012) or, as in the case of *lasR*, simply a reduction in the metabolic cost of maintaining an active regulatory response. Regardless of the specifics of the mechanism, our results suggest that mutations in *mexT* may evolve for reasons other than antibiotic selection. Others have observed the evolution of resistance in the absence of drug selection as well, and the genetic causes are often due to mutations in housekeeping genes like *rpoB* and *gyrA* (reviewed in Hershberg 2017). To the best of our knowledge, ours is the first study to show that *mexT* mutations can be selected in the absence of drug as well.

## Conclusions

Our work provides a snapshot of genomic changes associated with rapid adaptation and diversification in populations of *P. aeruginosa* evolving under conditions that mimic, to varying degrees, the nutritional complexity and viscosity of the CF lung environment. Our leading result is that genetic variation is abundant across all conditions including those resembling the CF lung and, unlike what was observed for phenotypic disparity (Schick and Kassen 2018), is not substantially higher in CF-like conditions. This result is attributable to high mutation supply rates resulting from large population sizes (∼10^8^ CFU/ml) and consistent with that seen in other microbial evolution experiments (Schick et al 2015; Good et al 2017). Because population densities in our experiment are, by design, similar to those in chronically-infected patients, this result suggests that *P. aeruginosa* populations in chronic infections can be highly diverse due to mutation alone. Genetic diversity may be higher still if distinct strains co-infect a single patient (Caballero et al 2015) and recombination can generate additional variants (Darch et al 2015).

We also saw low levels of genomic repeatability across all environments, a result in line with what is often seen in evolve-and-resequence experiments in bacteria and yeast (Bailey et al 2017). Importantly, however, evolution in CF-like conditions causes parallelism to be substantially lower than in the MIN environment, the least CF-like treatment in our experiment. This result is attributable both to divergent selection generated by nutritional complexity and spatial structure imposed by mucin. Divergent selection promotes the evolution of divergent niche specialists, reducing the likelihood of parallelism relative to directional selection towards a single fitness optimum. Mucin, for its part, reduces dispersal and creates spatially structured subpopulations with smaller effective population sizes than would be found under well-mixed conditions, making it less likely that the same beneficial mutations can be found and fixed by selection repeatedly in independently-evolved populations. This result mirrors what is seen *in vivo* among CF isolates (Marvig et al 2015, Williams et al 2015) and reinforces the notion that adaptation to the CF airway can follow many genetic routes, making it difficult to identify reliable genomic signals marking the transition from transient to chronic infection.

Despite the low levels of parallelism, on average, in our experiment, we did recover mutations in a number of genes thought to be important during adaptation to the CF airway. Many of these genes are likely to impact regulatory functions associated with quorum sensing or motility (*lasR, mvfR, wspF*) or motility itself (*wspA, pilA-D*). Importantly few, if any, of these mutations were specific to CF-like conditions and some, like *lasR*, even evolved repeatably across every environment in our experiment. These results suggest that many of the genetic changes observed in our experiment confer fitness benefits across a wide range of conditions rather than being adaptations specific to nutrient complexity or mucin in the CF lung. Indeed, changes to patterns of gene regulation, often mediated by loss-of-function mutations in non-essential genes, are commonly observed during the early stages of adaptation to novel, stressful environments in many selection experiments (Kassen 2014, Dettman et al 2012). Our results suggest the same might be true for the suite of genetic changes commonly recovered among *P. aeruginosa* isolates from CF patients. If so, then adaptation to the CF lung may be best understood as a particular instance of the more general phenomenon of adaptation to a stressful environment.

The strength of these inferences must, of course, be tempered by the fact that our experiment was done in the lab under conditions that are substantially different from those encountered in the CF lung. Most obviously, the populations we have studied here have evolved in the absence of an active immune system, competing microflora, or the regular administration of antibiotics. It is not immediately obvious how the quantity of genetic variation and degree of parallelism would respond to these additional sources of selection. On the one hand we might expect lower levels of genetic variation and higher parallelism if these stressors represent additional stressors that favour only those genotypes that can withstand the multiple sources of selection in the CF lung. Alternatively, if no single genotype is superior across all niche dimensions, then these additional stressors could serve to actively preserve genetic diversity in the CF airway. Nevertheless, it is clear that the large population sizes typical of *P. aeruginosa* infections can supply high levels of genetic variation through mutation alone, a necessary condition for adaptive evolution through selection. This interpretation lends support to the idea that the dynamics of genetic variation among *P. aeruginosa* in the CF lung can be caused mainly by mutation-driven selection.

## Materials and Methods

### Bacterial strains

Bacterial strains and populations used for this study were from the end-point of a selection experiment described in Schick and Kassen (2018). Briefly, a total of 120 populations derived from a common ancestor, *Pseudomonas aeruginosa* strain 14 (Pa14), were propagated in daily batch culture for ∼220 generations in one of four environments: SCFM (synthetic cystic fibrosis medium), MIN (M9 minimal salts + glucose), SCFM + mucin, and MIN + mucin. The factorial design of the experiment allows us to examine the main effects of nutritional complexity (SCFM versus MIN), spatial structure (mucin versus no mucin), and their interaction on phenotypic and genetic diversification. We randomly chose 24 populations from each of the SCFM and SCFM + mucin treatments, and 23 populations from each of the MIN and MIN + mucin treatments, for sequencing. Two reference strains, both ancestral to the selection experiment (Pa14 and a *lacZ*- marked Pa14, were also sequenced to facilitate genome assembly and to identify genetic variants evolved over the course of the experiment.

### Whole-genome sequencing

Populations were revived overnight from frozen in liquid Luria Bertani (LB) broth at 37°C. Genomic DNA was extracted for whole-genome sequencing from samples using the MO BIO Ultraclean 96 Microbial DNA kit (now sold as QIAGEN DNeasy UltraClean 96 Microbial kit), following the manufacturer’s recommended protocol. Library preparation and sequencing was performed by Genome Quebec at McGill University on the Illumina HiSeq 4000 platform, using paired-end sequencing of 2×100 base-pair reads.

### Sequence processing and generating variant table

Whole genome sequencing yielded a total of 75 Gb of raw data, with a median coverage of 120- fold. Analyses were performed in-house using a custom pipeline. Briefly, sequencing reads were quality trimmed using Trimmomatic version 0.36 (Bolger et al 2014), removing the leading and trailing bases below a quality score of 5 as well as scanning the read with a 5-base sliding window, cutting when the average quality per base drops below 20. Reads shorter than 20 base pairs were also discarded. Reads were then aligned to *Pseudomonas aeruginosa* reference genome UCBPP-PA14 109 (from Winsor et al 2016) using the bwa-mem algorithm of BWA version 0.7.12 and Samtools version 1.3.1 (Li et al 2009). Picard Tools version 2.9.2 was then used to mark PCR duplicates and add read group information. Variants were called using two independently developed algorithms. First, Breseq version 0.30.0 (Deatherage and Barrick 2014) was used with the predict polymorphism mode, a tool specifically designed for detecting mutations in microbial genomes. To confirm no major biases in variant calling, the HaplotypeCaller function of GATK version 3.6 (McKenna et al 2010) was used as well. To identify only mutations that arose over the course of the selection experiment, the set of variants found in our two ancestral strains (Pa14 and Pa14-*lacZ*) were compared to those found in our evolved populations. Variants in common across ancestral strains and all evolved populations were discarded. Of the 94 populations sequenced, we found 4 populations to have evidence of cross-contamination (1 pair each of SCFM and SCFM+mucin populations) and 9 populations to have increased mutation rates, evidenced by both a larger number of variants present (> 30) and variants in one of the following genes: *mutS* (2), *lexA, dnaA, dnaX, recQ, uvrD, ruvB*, and PA14_25780, all of which have been linked to increased mutation rates. These putative mutator variants were found in 4 SCFM populations, 3 SCFM+mucin populations, and 2 MIN+mucin populations. Both the populations with evidence of cross-contamination and populations with these putative hypermutators present were excluded from subsequent analysis.

### Statistical analysis

#### Estimating rates of genomic hitchhiking

To estimate the proportion of mutations that are selectively neutral but arose in a genome containing a beneficial mutation (a phenomenon referred to as hitchhiking), we compared the observed number of nonsynonymous mutations to the expected number of nonsynonymous mutations under neutrality, inferring that the excess mutations over the expected number are likely non-neutral, or ‘driver’ mutations. The expected number of nonsynonymous mutations under neutrality is based on the observed number of synonymous mutations, genetic changes assumed to have no effect on fitness.

Under neutral evolution, nonsynonymous and synonymous mutations arise at the same rate, accumulating to numbers proportional to the number of sites of each type. Using this estimate of the number of hitchhiker mutations (expected nonsynonymous mutations under neutrality), we determine the estimated number of driver mutations by subtracting the expected number of nonsynonymous mutations from the observed number of nonsynonymous mutations. We apply this calculation to all populations grouped together as well as populations within a given treatment. For all populations grouped together, we estimate that 28.4% of nonsynonymous SNPs are drivers (see Table 1 for mutation numbers). Further, after estimating the relative rates of hitchhiker and driver mutations, we compare the distribution of mutation frequencies of synonymous and nonsynonymous mutations to find that nearly all synonymous mutations are present at frequencies less than 0.16. Hitchhiker mutations are likely to be present at frequencies equal to or less than their respective driver mutations, suggesting that low frequency mutations are most likely to be hitchhikers. Therefore, we exclude low-frequency mutations to match the estimated proportion of hitchhiker mutation as calculated above. This corresponds to a frequency of 0.16, shown in Figure 2.

#### Genetic diversity

We estimated within-population genetic diversity by calculating heterozygosity, defined here as the mean heterozygosity (2pq, where p is the estimated allele frequency and q = 1-p) of all polymorphic loci in a population, excluding synonymous and low-frequency variants (as defined above). Treatment group ranks were preserved when low-frequency variants were included. Significant differences between treatment groups were tested for using a single-factor ANOVA. To quantify genetic divergence among evolved populations, we performed a principal coordinates analysis (PCoA) using the R package vegan (version 2.5.2) on a Euclidean distance matrix. We estimated genetic divergence within a treatment by calculating mean distance to the spatial median using the ‘betadisper’ function. This function determines if treatment groups differ in dispersion (variance), a multivariate analogue of the Levene test for homogeneity of variances. Following this, we performed a Tukey HSD test to determine which treatment groups differed significantly in mean dispersion. Dispersion was also used a measure of parallelism within treatment, discussed in the section below.

#### Parallelism and repeatability

We quantified population levels of parallelism using three different metrics: dispersion, Jaccard, and C-scores. All three metrics were calculated using the same set of data: frequencies of mutations in all genes, after excluding synonymous and low-frequency variants. For dispersion, we calculated the mean distance between a population and the treatment centroid, following a principal coordinates analysis (PCoA) on a Euclidean distance matrix. The measurement corresponds to treatment level genetic variance with larger mean dispersion signifying more divergent populations and therefore less parallelism. We also include standard error of dispersion. To determine significance, we performed an ANOVA with Euclidean distance as the response variable and treatment as the explanatory. For the Jaccard measure, we used the Jaccard index to calculate the dissimilarity between all pairs of populations within a treatment. We then report the mean and standard error of all pairwise comparisons, with larger values signifying larger dissimilarities and therefore less parallelism. C-score is a metric for repeatability that uses the hypergeometric distribution to calculate the deviation between the observed amount of parallelism and the expectation under random gene-use (Yeaman et al 2018). The magnitude of the C-score represents the magnitude of the deviation with larger C-scores signify higher repeatability and therefore more parallelism. To determine significance for both Jaccard and C-score metrics, we performed an exact test (permutation test) by randomizing treatment labels (number of permutations = 10000) and calculating a null distribution of F-values.

Parallelism at the gene level was defined as the proportion of populations with mutations in that gene, both globally for all treatments and within specific treatments. To test for significance, we calculated the probability of our observed results if gene use was random, using the binomial distribution with the number of populations as the number of trials, number of times a gene was mutated as the number of successes, and proportion of total populations across all treatments with a mutation in that gene as the probability of success. From this, if the probability of an observation was less than 0.05, we considered that gene to be treatment specific.

## Acknowledgments

This work was supported by a Natural Sciences and Engineering Research Council (Canada) Discovery Grant to RK.

## Supplementary Figures

**Supplementary Figure 1.**
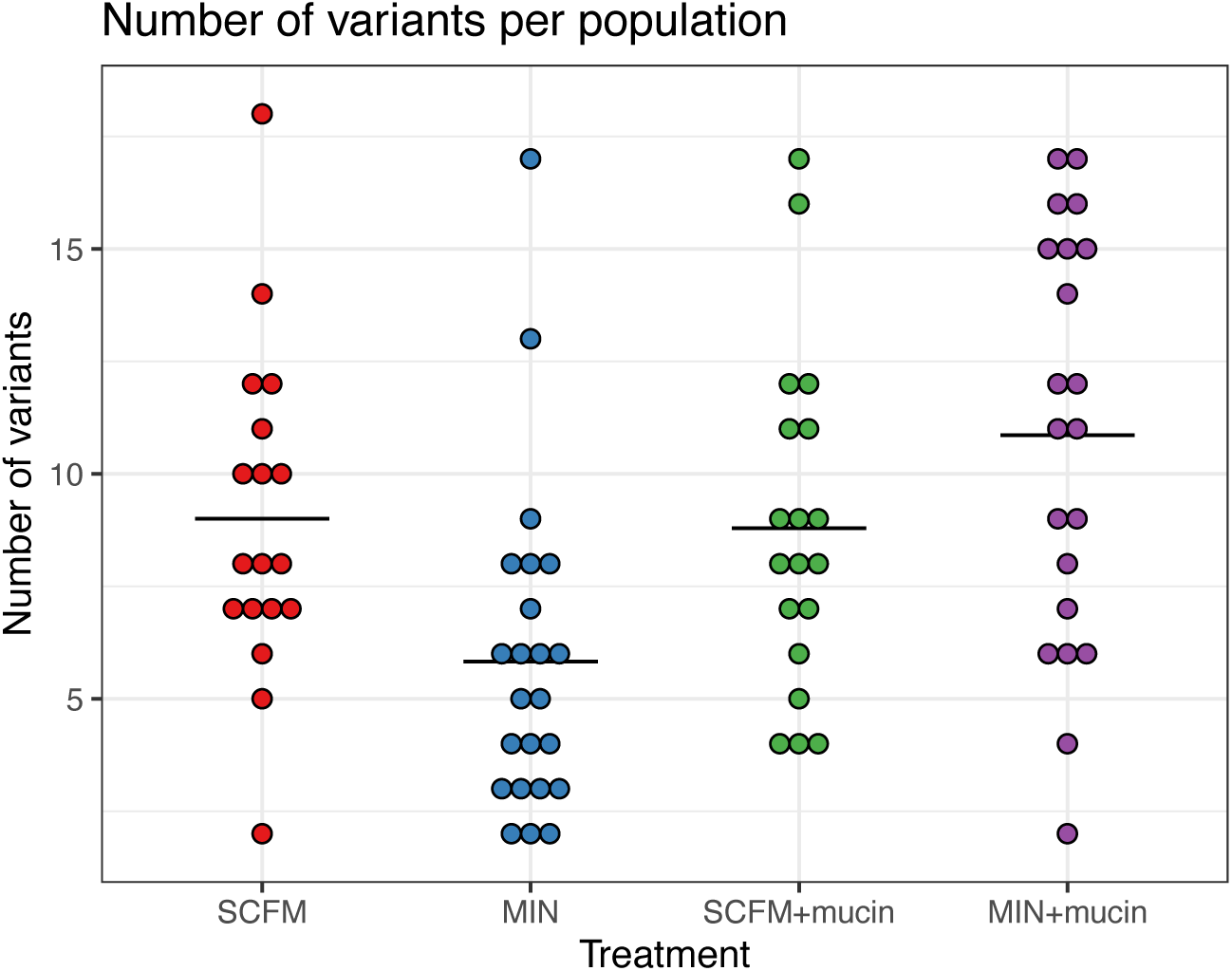
Number of variants per population. Number of variants discovered in each population, including low frequency SNPs. Horizontal bars represent treatment means.

**Supplementary Figure 2.**
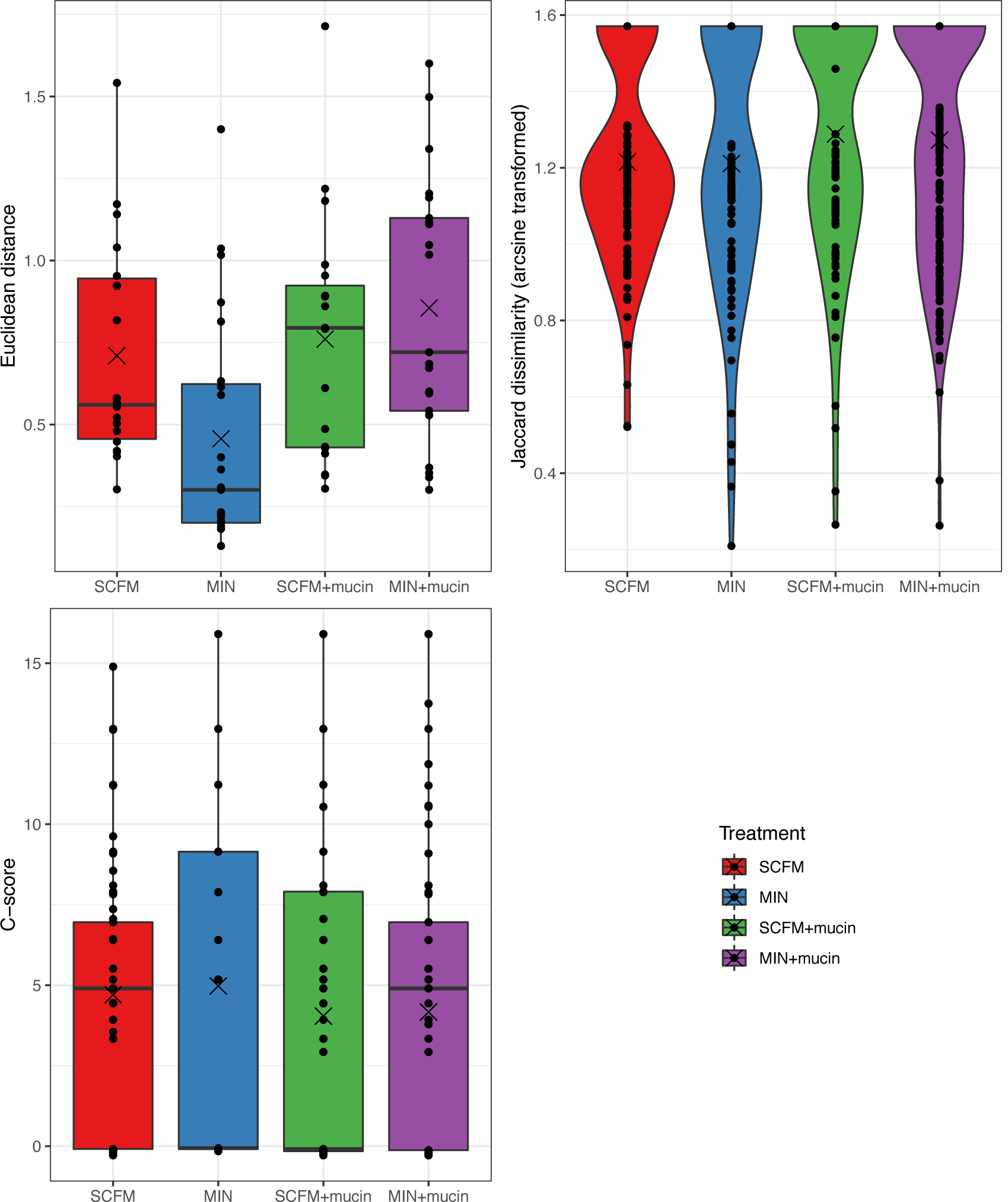
Genetic divergence among populations. **A**. Euclidean distance is the distance each population is from the centroid of the treatment group, based on a principal coordinates analysis (PCoA). **B**. Jaccard dissimilarity, arcsine transformed for visualization, of all pairwise comparisons between populations within a treatment group. **C**. C-scores, magnitude of genetic constraint, for all pairwise comparisons between populations within a treatment group. In all panels, horizontal bars denote treatment median and crosses denote treatment mean.

**Supplementary Figure 3.**
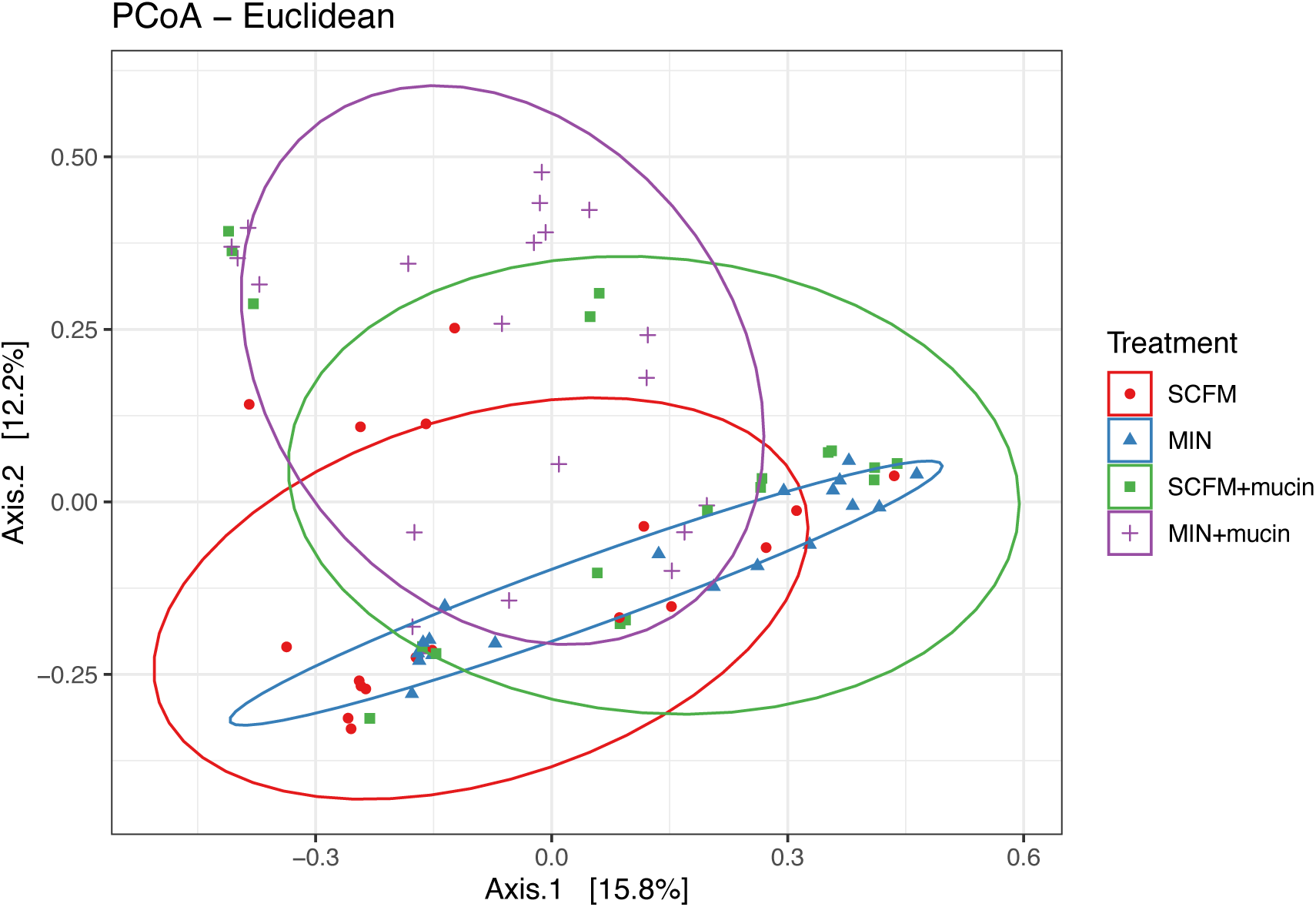
Diversity among populations. The first two axes of a principal coordinates analysis (PCoA), based on a Euclidean distance matrix. Each point represents a population, and ellipses represent a 90 percent confidence interval around a multivariate t- distribution.

**Supplementary Figure 4.**
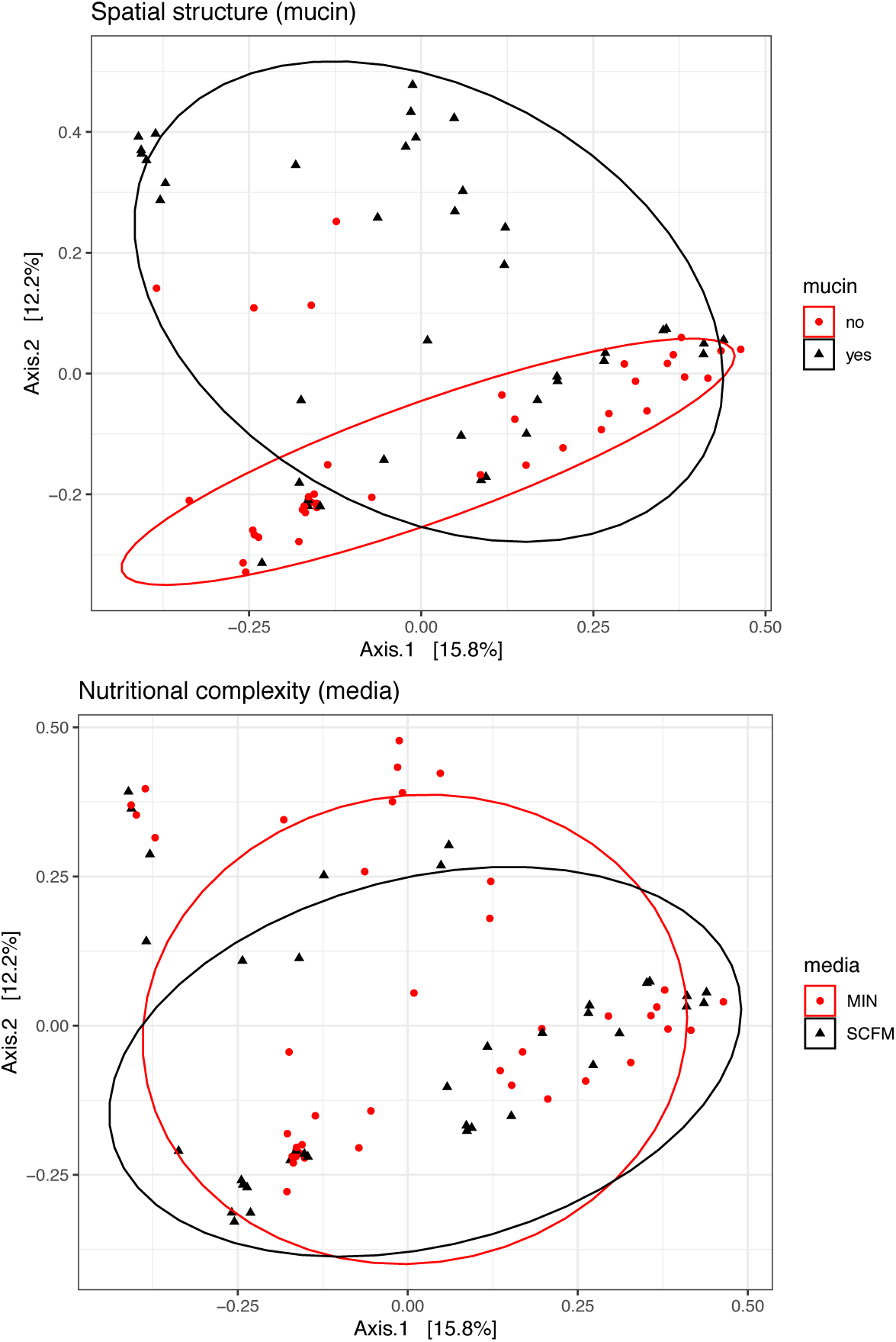
Diversity among populations with effects of media and mucin separately. The first two axes of a principal coordinates analysis (PCoA), based on a Euclidean distance matrix. Populations are classified by media (SCFM or MIN; A), and mucin (presence or absence; B). Each point represents a population, and ellipses represent a 90 percent confidence interval around a multivariate t-distribution.

## References

Agashe D, Sane M, Phalnikar K, Diwan GD, Habibullah A, Martinez-Gomez NC, Sahasrabuddhe V, Polachek W, Wang J, Chubiz LM, Marx CJ. 2016. Large-Effect Beneficial Synonymous Mutations Mediate Rapid and Parallel Adaptation in a Bacterium. Mol Biol Evol. 33:1542–1553.

Ashish A, Paterson S, Mowat E, Fothergill JL, Walshaw MJ, Winstanley C. 2013. Extensive diversification is a common feature of Pseudomonas aeruginosa populations during respiratory infections in cystic fibrosis. Journal of Cystic Fibrosis. 12:790–793.

Bailey SF, Blanquart F, Bataillon T, Kassen R. 2017. What drives parallel evolution : How population size and mutational variation contribute to repeated evolution. BioEssays. 39:e201600176.

Bailey SF, Rodrigue N, Kassen R. 2015. The effect of selection environment on the probability of parallel evolution. Mol Biol Evol 32:1436–1448.

Bailey SF, Hinz A, Kassen R. 2014. Adaptive synonymous mutations in an experimentally evolved Pseudomonas fluorescens population. Nature Communications [Internet]. [cited 2018 Mar 19]; 5.

Behringer MG, Choi BI, Miller SF, Doak TG, Karty JA, Guo W, Lynch M. 2018. *Escherichia coli* cultures maintain stable subpopulation structure during long-term evolution. Proceedings of the National Academy of Sciences. 115:E4642–E4650.

Bolger AM, Lohse M, Usadel B. 2014. Trimmomatic: a flexible trimmer for Illumina sequence data. Bioinformatics. 30:2114–2120.

Breidenstein EBM, de la Fuente-Núñez C, Hancock REW. 2011. Pseudomonas aeruginosa: all roads lead to resistance. Trends in Microbiology. 19:419–426.

Burrows LL. 2012. *Pseudomonas aeruginosa* Twitching Motility: Type IV Pili in Action. Annu Rev Microbiol. 66:493–520.

Buskirk SW, Peace RE, Lang GI. 2017. Hitchhiking and epistasis give rise to cohort dynamics in adapting populations. Proceedings of the National Academy of Sciences. 114:8330–8335.

Darch SE, McNally A, Harrison F, Corander J, Barr HL, Paszkiewicz K, Holden S, Fogarty A, Crusz SA, Diggle SP. 2015. Recombination is a key driver of genomic and phenotypic diversity in a Pseudomonas aeruginosa population during cystic fibrosis infection. Scientific Reports [Internet]. [cited 2017 Oct 10]; 5

D’Argenio DA, Wu M, Hoffman LR, Kulasekara HD, Déziel E, Smith EE, Nguyen H, Ernst RK, Larson Freeman TJ, Spencer DH, Brittnacher M. 2007. Growth phenotypes of Pseudomonas aeruginosa lasR mutants adapted to the airways of cystic fibrosis patients. Molecular microbiology. Apr;64(2):512–33.

Deatherage DE, Barrick JE. 2014. Identification of Mutations in Laboratory-Evolved Microbes from Next-Generation Sequencing Data Using breseq. In: Sun L, Shou W, editors. Engineering and Analyzing Multicellular Systems [Internet]. Vol. 1151. New York, NY: Springer New York; [cited 2018 Jul 26]; p. 165–188.

Dettman JR, Rodrigue N, Melnyk AH, Wong A, Bailey SF, Kassen R. 2012. Evolutionary insight from whole-genome sequencing of experimentally evolved microbes. Molecular Ecology. 21:2058–2077.

Dettman JR, Rodrigue N, Aaron SD, Kassen R. 2013. Evolutionary genomics of epidemic and nonepidemic strains of *Pseudomonas aeruginosa*. Proc Natl Acad Sci USA 110:21065–21070.

Diaz Caballero J, Clark ST, Coburn B, Zhang Y, Wang PW, Donaldson SL, Tullis DE, Yau YCW, Waters VJ, Hwang DM, Guttman DS. 2015. Selective Sweeps and Parallel Pathoadaptation Drive *Pseudomonas aeruginosa* Evolution in the Cystic Fibrosis Lung. mBio. 6:e00981–15.

Fargier E, Mac Aogain M, Mooij MJ, Woods DF, Morrissey JP, Dobson ADW, Adams C, O’Gara F. 2012. MexT Functions as a Redox-Responsive Regulator Modulating Disulfide Stress Resistance in Pseudomonas aeruginosa. Journal of Bacteriology. 194:3502–3511.

Feltner JB, Wolter DJ, Pope CE, Groleau M-C, Smalley NE, Greenberg EP, Mayer-Hamblett N, Burns J, Déziel E, Hoffman LR, Dandekar AA. 2016. LasR Variant Cystic Fibrosis Isolates Reveal an Adaptable Quorum-Sensing Hierarchy in *Pseudomonas aeruginosa*. mBio [Internet]. [cited 2018 Jul 26]; 7.

Flynn KM, Dowell G, Johnson TM, Koestler BJ, Waters CM, Cooper VS. 2016. Evolution of ecological diversity in biofilms of *Pseudomonas aeruginosa* by altered cyclic diguanylate signaling. J Bacteriol 198: 2608–2618.

Foweraker JE, Laughton CR, Brown DF, Bilton D. 2005. Phenotypic variability of Pseudomonas aeruginosa in sputa from patients with acute infective exacerbation of cystic fibrosis and its impact on the validity of antimicrobial susceptibility testing. Journal of Antimicrobial Chemotherapy. 55(6):921–7.

Freschi L, Bertelli C, Jeukens J, Moore MP, Kukavica-Ibrulj I, Emond-Rheault J-G, Hamel J, Fothergill JL, Tucker NP, McClean S, et al. 2018. Genomic characterisation of an international Pseudomonas aeruginosa reference panel indicates that the two major groups draw upon distinct mobile gene pools. FEMS Microbiology Letters [Internet]. [cited 2018 Oct 4]; 365.

Futuyma DJ, Moreno G. 1988. The evolution of ecological specialization. Annual Review of Ecology and Systematics. 19:207–233.

Good BH, McDonald MJ, Barrick JE, Lenski RE, Desai MM. 2017. The dynamics of molecular evolution over 60,000 generations. Nature. 551:45–50.

Guillaume Martin, Thomas Lenormand. 2006. A General Multivariate Extension of Fisher’s Geometrical Model and the Distribution of Mutation Fitness Effects across Species. Evolution. 60:893–907.

Hershberg R. 2017. Antibiotic-Independent Adaptive Effects of Antibiotic Resistance Mutations. Trends in Genetics. 33:521–528.

Hickman JW, Tifrea DF, Harwood CS. 2005. A chemosensory system that regulates biofilm formation through modulation of cyclic diguanylate levels. Proc Natl Acad Sci USA 102: 14422–14427.

Hoffman LR, Kulasekara HD, Emerson J, Houston LS, Burns JL, Ramsey BW, Miller SI. 2009. Pseudomonas aeruginosa lasR mutants are associated with cystic fibrosis lung disease progression. Journal of Cystic Fibrosis. 8:66–70.

Hoiby N, Pressler T. 2006. Emerging pathogens in cystic fibrosis. European Respiratory Monograph. 35:66.

LaFayette SL, Houle D, Beaudoin T, Wojewodka G, Radzioch D, Hoffman LR, Burns JL, Dandekar AA, Smalley NE, Chandler JR, Zlosnik JE. 2015. Cystic fibrosis–adapted Pseudomonas aeruginosa quorum sensing lasR mutants cause hyperinflammatory responses. Science advances. 1;1(6):e1500199.

Kassen R. 2009. Toward a General Theory of Adaptive Radiation: Insights from Microbial Experimental Evolution. Annals of the New York Academy of Sciences. 1168:3–22.

Kassen R. 2014. Experimental evolution and the nature of biodiversity. Roberts.

Khan AI, Dinh DM, Schneider D, Lenski RE, Cooper TF. 2011. Negative Epistasis Between Beneficial Mutations in an Evolving Bacterial Population. Science. 332:1193–1196.

Kohler T, Epp SF, Curty LK, Pechere JC. 1999. Characterization of MexT, the regulator of the MexE-MexF-OprN multidrug efflux system of *Pseudomonas aeruginosa*. J Bacter 181:6300–6305.

Kristofich J, Morgenthaler AB, Kinney WR, Ebmeier CC, Snyder DJ, Old WM, Cooper VS, Copley SD. 2018. Synonymous mutations make dramatic contributions to fitness when growth is limited by a weak-link enzyme. PLoS genetics. 14(8):e1007615.

LaFayette SL, Houle D, Beaudoin T, Wojewodka G, Radzioch D, Hoffman LR, Burns JL, Dandekar AA, Smalley NE, Chandler JR, et al. 2015. Cystic fibrosis-adapted Pseudomonas aeruginosa quorum sensing lasR mutants cause hyperinflammatory responses. Science Advances. 1:e1500199–e1500199.

Leale AM, Kassen R. 2018. The emergence, maintenance, and demise of diversity in a spatially variable antibiotic regime. Evolution Letters. 2:134–143.

Lebeuf-Taylor E, McCloskey N, Bailey SF, Hinz A, Kassen R. 2019. The distribution of fitness effects among synonymous mutations in a gene under directional selection. eLife. 8:e45952.

Li H, Handsaker B, Wysoker A, Fennell T, Ruan J, Homer N, Marth G, Abecasis G, Durbin R, 1000 Genome Project Data Processing Subgroup. 2009. The Sequence Alignment/Map format and SAMtools. Bioinformatics. 25:2078–2079.

Markussen T, Marvig RL, Gomez-Lozano M, Aanaes K, Burleigh AE, Hoiby N, Johansen HK, Molin S, Jelsbak L. 2014. Environmental Heterogeneity Drives Within-Host Diversification and Evolution of Pseudomonas aeruginosa. mBio. 5:e01592-14-e01592-14.

Marvig RL, Sommer LM, Molin S, Johansen HK. 2015. Convergent evolution and adaptation of Pseudomonas aeruginosa within patients with cystic fibrosis. Nature Genetics. 47:57–64.

McKenna A, Hanna M, Banks E, Sivachenko A, Cibulskis K, Kernytsky A, Garimella K, Altshuler D, Gabriel S, Daly M, DePristo MA. 2010. The Genome Analysis Toolkit: A MapReduce framework for analyzing next-generation DNA sequencing data. Genome Research. 20:1297–1303.

Mowat E, Paterson S, Fothergill JL, Wright EA, Ledson MJ, Walshaw MJ, Brockhurst MA, Winstanley C. 2011. *Pseudomonas aeruginosa* Population Diversity and Turnover in Cystic Fibrosis Chronic Infections. American Journal of Respiratory and Critical Care Medicine. 183:1674–1679.

Poltak SR, Cooper VS. 2011. Ecological succession in long-term experimentally evolved biofilms produces synergistic communities. ISME 5:369–378.

Poole K. 2005. Efflux-mediated antimicrobial resistance. Journal of Antimicrobial Chemotherapy. 56:20–51.

Rajan S, Saiman L. 2002. Pulmonary infections in patients with cystic fibrosis. Seminars in respiratory infections. Vol. 17, No. 1, pp. 47–56.

Schaedel C, de Monestrol I, Hjelte L, Johannesson M, Kornfält R, Lindblad A, Strandvik B, Wahlgren L, Holmberg L. 2002. Predictors of deterioration of lung function in cystic fibrosis*: CFTR Genotype in Lung Function in CF. Pediatric Pulmonology. 33:483–491.

Schick A, Bailey SF, Kassen R. 2015. Evolution of Fitness Trade-Offs in Locally Adapted Populations of *Pseudomonas fluorescens*. The American Naturalist. 186:S48–S59.

Schick A, Kassen R. 2018. Rapid diversification of *Pseudomonas aeruginosa* in cystic fibrosis lung-like conditions. Proc Natl Acad Sci USA. 115:10714–10719.

Schluter. 2000. Ecological Character Displacement in Adaptive Radiation. The American Naturalist. 156:S4.

Smith EE, Buckley DG, Wu Z, Saenphimmachak C, Hoffman LR, D’Argenio DA, Miller SI, Ramsey BW, Speert DP, Moskowitz SM, others. 2006. Genetic adaptation by Pseudomonas aeruginosa to the airways of cystic fibrosis patients. Proceedings of the National Academy of Sciences. 103:8487–8492.

Stressman FA, Rogers GB, Marsh P, et al. 2011. Does bacterial density in cystic fibrosis sputum increase prior to pulmonary exacerbation? J Cystic Fibrosis 10:357–365.

Traverse CC, Mayo-Smith LM, Poltak SR, Cooper VS. 2013. Tangled bank of experimentally evolved *Burkholderia* biofilms reflects selection during chronic infections. Proc Natl Acad Sci 110: E250–E259.

Turner CB, Marshall CW, Cooper VS. 2018. Parallel genetic adaptation across environments differing in mode of growth or resource availability. Evolution Letters 2-4: 355–367.

Williams D, Evans B, Haldenby S, Walshaw MJ, Brockhurst MA, Winstanley C, Paterson S. 2015. Divergent, coexisting *Pseudomonas aeruginosa* lineages in chronic cystic fibrosis lung infections. Am J Respir Crit Care Med 191:775–785.

Winsor GL, Griffiths EJ, Lo R, Dhillon BK, Shay JA, Brinkman FSL. 2016. Enhanced annotations and features for comparing thousands of *Pseudomonas* genomes in the Pseudomonas genome database. Nucleic Acids Research. 44:D646–D653.

Workentine ML, Sibley CD, Glezerson B, Purighalla S, Norgaard-Gron JC, Parkins MD, Rabin HR, Surette MG. 2013. Phenotypic Heterogeneity of Pseudomonas aeruginosa Populations in a Cystic Fibrosis Patient. Battista JR, editor. PLoS ONE. 8:e60225.

Wright EA, Fothergill JL, Paterson S, Brockhurst MA, Winstanley C. 2013. Sub-inhibitory concentrations of some antibiotics can drive diversification of Pseudomonas aeruginosa populations in artificial sputum medium. BMC microbiology. 13:170.

Yang L, Jelsbak L, Marvig RL, Damkiaer S, Workman CT, Rau MH, Hansen SK, Folkesson A, Johansen HK, Ciofu O, et al. 2011. Evolutionary dynamics of bacteria in a human host environment. Proceedings of the National Academy of Sciences. 108:7481–7486.

Yeaman S, Gerstein AC, Hodgins KA, Whitlock MC. Quantifying how constraints limit the diversity of viable routes to adaptation. :25.

